# Arabidopsis phytochelatin synthase 1, but not phytochelatin synthesis, functions in extracellular defense against multiple fungal pathogens

**DOI:** 10.1101/568113

**Authors:** Kian Hematy, Melisa Lim, Candice Cherk, Paweł Bednarek, Mariola Piślewska-Bednarek, Clara Sanchez-Rodriguez, Monica Stein, Rene Fuchs, Christine Klapprodt, Volker Lipka, Antonio Molina, Erwin Grill, Paul Schulze-Lefert, Shauna Somerville

## Abstract

Phytochelatin synthase (PCS) is a key component of heavy metal detoxification in plants. PCS catalyzes both the synthesis of the peptide phytochelatin from glutathione as well as the degradation of glutathione conjugates via peptidase activity. Here, we describe a hitherto uncharacterized role for PCS in disease resistance against plant pathogenic fungi. The *pen4* mutant, which is allelic to *cadmium insensitive 1* (*cad1/pcs1*) mutants, was recovered from a screen for Arabidopsis mutants with reduced resistance to the non-adapted barley fungal pathogen, *Blumeria graminis* f. sp. *hordei*. PCS1, which is found in the cytoplasm of cells of healthy plants, translocates upon pathogen attack and colocalizes with the PEN2 myrosinase on the surface of immobilized mitochondria. *pcs1* and *pen2* mutant plants exhibit a similar metabolic defect in the accumulation of pathogen-inducible indole glucosinolate-derived compounds, suggesting that PEN2 and PCS1 act in the same metabolic pathway. The function of PCS1 in this pathway is independent of phytochelatin synthesis and deglycination of glutathione conjugates, as catalytic-site mutants of PCS1 are still functional in indole glucosinolate metabolism. In uncovering a previously unknown function for PCS1, we reveal this enzyme to be a moonlighting protein important for plant responses to both biotic and abiotic stresses.

---

As sessile organisms, plants have developed sophisticated mechanisms that allow them to adapt to and survive the biotic and abiotic stresses that they encounter in their environments. Different stress signaling pathways are connected to maximize plant fitness (Berens et al., 2019). For instance, one evolutionary conserved mechanism involves phytohormone signaling mediated by abscisic acid, which promotes abiotic stress tolerance and suppresses signaling of the biotic stress-related phytohormone salicylic acid (Berens et al., 2017; Berens et al., 2019). On the other hand, an overlapping enzymatic machinery is engaged to cope with xenobiotics and pathogen stress. The metabolism of xenobiotics, including plant metal tolerance, as well as the biosynthesis of sulfur-containing secondary metabolites that act as antimicrobials and/or activate evolutionarily conserved immune responses, involves in Brassicaceae species the formation and processing of glutathione conjugates (Grill et al., 1989; Cobbett and Goldsbrough, 2002; Bednarek et al., 2009; Pastorczyk and Bednarek, 2016; Czerniawski and Bednarek, 2018).

Invasive growth is an essential part of the pathogenesis of many fungal and oomycete plant pathogens, and extracellular defense mechanisms play a critical role in blocking their entry into plant cells (Underwood and Somerville, 2008; Hematy et al., 2009). For example, *Arabidopsis thaliana* is an inappropriate or non-host plant for the non-adapted barley powdery mildew pathogen, *Blumeria graminis sp. hordei* (*Bgh*) (Lipka et al., 2008). The ability of Arabidopsis cells to resist *Bgh* entry is partly dependent on the PENETRATION (PEN) proteins. For instance, mutants of the syntaxin PEN1 (=SYP121) (Collins et al., 2003), the myrosinase PEN2 (Lipka et al., 2005) and the ABC transporter PEN3 (=PDR8) (Stein et al., 2006) are more susceptible to *Bgh* entry and exhibit increased formation of the fungal feeding structure, the haustoria, in leaf epidermal cells. It has been shown that PEN2 hydrolyzes indole glucosinolates (IGs), a class of sulfur-containing secondary metabolites, to form products that could act as broad spectrum toxins and confer antifungal defense in the extracellular space at pathogen contact sites (Bednarek et al., 2009). It has also been suggested that IG-derived compounds are signaling molecules for callose deposition following treatment with the microbe-associated molecular pattern (MAMP) flagellin (flg22), a peptide epitope derived from the bacterial motor protein (Clay et al., 2009). Recently, Arabidopsis Glutathione-*S*-Transferase class-tau member 13 (GSTU13) was identified as an indispensable component of the PEN2 immune pathway for IG metabolism (Piślewska-Bednarek et al., 2018), suggesting that this pathway involves conjugation of the tripeptide glutathione (ECG) with unstable isothiocyanates (ITCs) that are products of IG metabolism and further processing of the resulting adducts to biologically active molecules. Arabidopsis PAD2/CAD2, which encodes γ-glutamylcysteine synthetase (γ-ECS) and catalyzes the first committed step of glutathione biosynthesis, is essential for biotic stress-induced accumulation of IGs and the PEN2 immune pathway (Bednarek et al., 2009). The same enzyme is also needed for heavy metal tolerance, as glutathione is the precursor of heavy-metal chelating polypeptides, called phytochelatins (Grill et al., 1985; Grill et al., 1989; Howden et al., 1995).

We present here the characterization of PEN4, a new player in penetration resistance of Arabidopsis against powdery mildews. PEN4 corresponds to PCS1 (Grill et al., 1989; Clemens et al., 1999), which encodes phytochelatin synthase 1 (Rea et al., 2004), an enzyme that catalyzes the terminal step in the synthesis of the heavy metal-chelating polypeptide phytochelatin (Grill et al., 1985; Grill et al., 1989). Like PEN2, PCS1 appears to be involved in IG metabolism. Both *pcs1* and *pen2* mutants are hypersusceptible to non-adapted and host-adapted pathogenic fungi and fail to accumulate IG-derivatives after inoculation. Like other PEN proteins, PCS1 is translocated underneath pathogen contact sites, where it colocalizes with PEN2, which is known to accumulate on the surface of immobilized mitochondria (Fuchs et al., 2016). Mutagenesis of PCS1 residues abolishing phytochelatin synthesis affects tolerance to heavy metal but does not alter PCS1 function in IG metabolism or extracellular defense. Together our results show that PCS1 is a multi-functional protein with two independent activities. As described previously, this protein catalyzes the synthesis of the peptide phytochelatin for heavy-metal tolerance but it also plays a newly described role in IG metabolism and immune responses.

## RESULTS

### Mutations in *PHYTOCHELATIN SYNTHASE 1* result in enhanced invasive growth of several fungal pathogens

The *pen4* mutant was isolated in a screen for Arabidopsis lines that allowed increased penetration of the non-adapted barley powdery mildew *Bgh* into leaf cells (Figure 1) (Stein et al., 2006). Typically, 5–10% of germinated *Bgh* conidiospores successfully breach the plant cell wall and differentiate into fungal haustoria for nutrient uptake in Arabidopsis Col-0 (wild-type) leaf epidermal cells (Figure 1A, arrowhead), while the remaining attempts at fungal entry culminate in *de novo*-synthesized callose-rich deposits called papillae (Figure 1A, star). The frequency of haustorium formation by *Bgh* increases to 20–25% on the leaves of *pen4* plants (Figure 1C).

**Figure 1.**
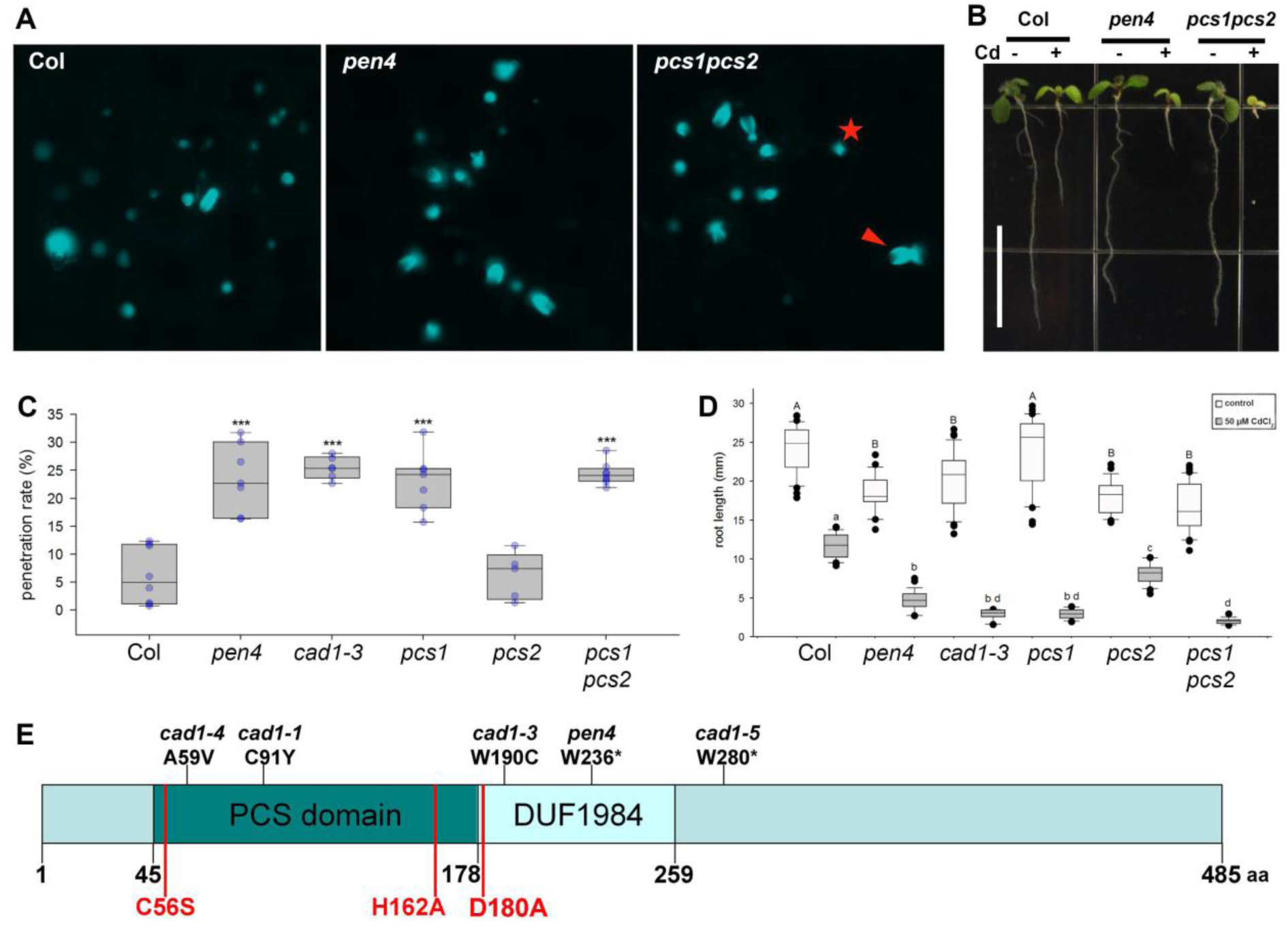
PEN4/PCS1 is important for both non-host penetration resistance and heavy metal tolerance. **(A)** Aniline Blue-stained leaves of 3-week-old Arabidopsis plants 24 h after infection by *Bgh*. Asterisk marks callose papillae symptomatic of failed penetration attempts, arrow head indicates stained callose-containing encasements of haustorial necks formed after a successful penetration by the fungal appressorium. **(B)** 7-day-old Arabidopsis seedlings of the indicated genotype grown in the presence or absence of cadmium (50 µM CdCl_2_). **(C)** Box plots showing penetration rate of *Bgh* spores on leaves of 3-week-old Arabidopsis plants. Measurements were done 24 h post-infection on the first true leaves of 5–6 independent plants (600–800 total penetration attempts counted per genotype). *** p<0.001 according to Student’s t-test. **(D)** Root length of 7-day-old seedlings grown on with or without 50 µM CdCl_2_. Different letters show statistical differences according to Tukey’s post-hoc test following one-way ANOVA. **(E)** Domain organization of PCS1 showing the position of several mutant alleles (above) and residues important for phytochelatin synthesis (below in red).

Map-based cloning of *PEN4* revealed that it encodes phytochelatin synthase (PCS1). PCS proteins are known for their glutathione γ-glutamylcysteinyltransferase activity (EC 2.3.2.15). The mutation in *pen4* results in a premature stop codon that truncates the protein at amino acid residue 236 (Figure 1E) in the middle of a DUF1984 domain (PF09328), which is always associated with the phytochelatin domain (PF05023) (Rea, 2006). To confirm that the mutation in *PCS1* is responsible for enhanced *Bgh* entry into *pen4* leaves, we tested another *pcs1* allele previously isolated in screens for plants with reduced tolerance to heavy metal cadmium ions (*cad1-3*) (Howden et al., 1995) and a null line carrying a T-DNA insertion in the second exon of *PCS1* (Blum et al., 2007). All *pcs1* lines were more susceptible to *Bgh* haustorium formation than the wild type Col-0 (Figure 1C). Like the parental lines, the F1 progeny of a cross between the *cad1-3* and *pen4* plants led to an increase of *Bgh* entry rates up to 20%, indicating that *pen4* is allelic to *cad1-3* (Figure S1).

Consistent with an impaired function of PCS1, the *pen4* mutant also showed decreased tolerance to cadmium (Figure 1B,D). Furthermore, the disease resistance defect of the *pen4* or *pcs1 pcs2* double mutant line could be complemented by transforming these mutants with *PCS1* expressed under the control of its native promoter or expressed under a 35S promoter and tagged with GFP at its N-terminus. Together, these data indicate that PCS1 plays a role in limiting plant cell entry by *Bgh*.

To determine whether PCS1 is important for resistance responses to other fungal pathogens, we inoculated *pen4* plants with the necrotrophic fungi *Botrytis cinerea* and *Plectosphaerella cucumerina* (Figure S2) (Sanchez-Rodriguez et al., 2009). Infection with both fungi was more severe in these mutant plants. Furthermore, *pen4* mutants, like *pen2* mutants, were hypersusceptible to powdery mildew adapted for growth on *A. thaliana*, *Golovinomyces cichoracearum* (Figure 2C). Together, these results show that PCS1 is important both in preventing the entry of non-adapted pathogens and in basal defense responses against multiple fungal pathogens.

**Figure 2.**
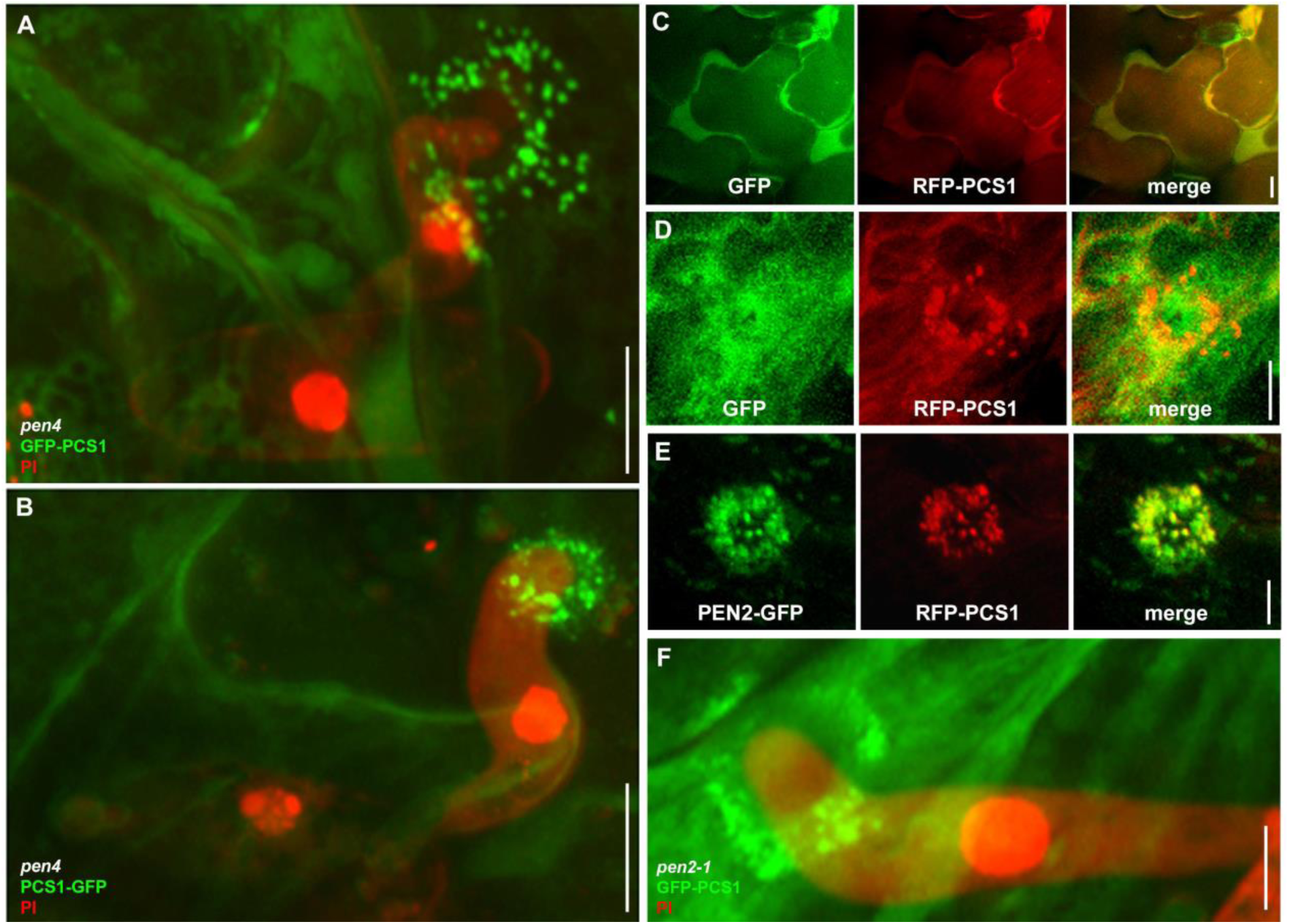
Pathogen attack triggers translocation of cytoplasmic PCS1 and mitochondrial colocalization with PEN2 at fungal penetration sites. 3-week-old Arabidopsis seedlings were infected with the non-adapted powdery mildew *Blumeria graminis sp. hordei* (*Bgh*). The epidermis of the first true leaves was visualized by confocal laser scanning microscopy 12–16 h after *Bgh* inoculation. The fungal structures are stained with propidium iodide except in GFP/RFP localization experiments. All pictures represent averaged Z-projection of stacks of 30–50 sections, except in **(C)** where a single median section is shown. Localization at the attempted penetration site of GFP-PCS1 **(A)** or PCS1-GFP **(B)** in a *pen4* mutant leaf epidermal cell. **(C)** Cytosolic localization of soluble GFP with RFP-PCS1 in non-infected epidermal cells. **(D)** Translocation of RFP-PCS1 but not of soluble GFP into aggregates at the penetration site. **(E)** Colocalization of RFP-PCS1 with mitochondria-associated PEN2-GFP at the penetration site. **(F)** Localization of GFP-PCS1 in a *pen2* mutant background after *Bgh* infection. Scale bar is either 10 µm (A-C) or 5 µm (D-F).

PCS2, the second isoform of the phytochelatin synthase in Arabidopsis, exhibits 83% overall amino acid identity with PCS1. Although the PCS2 protein is functional in synthesizing phytochelatins in response to heavy metals, the gene appears dispensable for heavy metal tolerance (Figure 1D, (Blum et al., 2007). We asked whether PCS2, like PCS1, had a role in penetration resistance to fungal pathogens. A mutant line harboring a T-DNA insertion that abolishes *PCS2* expression (*pcs2*) (Blum et al., 2007) allowed the same frequency of cell entry by *Bgh* as wild-type Col-0 plants (Figure 1C). Furthermore, a line harboring T-DNA insertions in both *PCS1* and *PCS2* (*pcs1 pcs2*) (Blum et al., 2007) did not consistently support a higher frequency of cell entry by *Bgh* than the *pcs1* single mutant (Figure 1C). As with heavy metal tolerance, PCS2 has no detectable role in extracellular defense to the non-adapted pathogen.

### Translocation of cytosolic PCS1 to immobile mitochondria underneath pathogen contact sites

Powdery mildew attack induces dramatic changes in cellular organization at the site of attempted fungal penetration (Underwood and Somerville, 2008). Several proteins that function in preventing host cell entry by *Bgh* show focal accumulation beneath fungal appressoria within epidermal cells at the site of attack, suggesting that these proteins act locally in extracellular defense responses (Koh et al., 2005; Underwood and Somerville, 2008). For instance, entry attempts of the fungus trigger local arrest of a subpopulation of mitochondria underneath *Bgh* appressoria, which is accompanied by transient aggregate formation of tail-anchored PEN2-GFP on the surface of these organelles (Fuchs et al., 2016). The plasma membrane-resident PEN3-GFP focally accumulates as disks, sometimes with bullseye-like “rings” around these contact sites (Koh et al., 2005; Stein et al., 2006).

Given PCS1’s involvement in limiting host cell entry, we tested whether PCS1 localized to the site of fungal attack. To this end, we generated GFP fusions at either the amino- or carboxy-terminus of PCS1 under control of the 35S promoter in the *pen4* background (Figure 2A, B). In addition, we generated a transgenic *pcs1 pcs2* double mutant line expressing RFP-PCS1 driven by the 35S promoter. In non-inoculated leaf epidermal cells, the RFP-PCS1 fluorescence signal suggested a cytosolic localization similar to that of free GFP expressed under the control of the 35S promoter (EGAD line; (Cutler et al., 2000); Figure 2C). This is consistent with the absence of canonical organelle targeting sequences or predicted transmembrane domains in PCS1. In pathogen-challenged leaf epidermal cells, in addition to their cytosolic localization, GFP-PCS1 and PCS1-GFP localized to bright round aggregates underneath *Bgh* contact sites (Figure 2A, B); GFP alone did not exhibit this localization (Figure 2D), excluding the possibility that the fluorescent tag is itself driving PCS1 aggregation underneath pathogen contact sites. Similarly, RFP-PCS1 also accumulated focally at incipient *Bgh* entry sites (Figure 2D, E). During the early stages of fungal entry, PCS1 tightly localized just beneath the tip region of the appressorium (Figure 2B,E), while at later stages the aggregates appeared more dispersed and excluded from the center of the penetration site (Figure 2A, D, F). Furthermore, the signal recruiting PCS1 to attempted entry sites was able to cross cell frontiers (Figure 2F, GFP-PCS1 from two adjacent cells organized as a ring around the fungal penetration site).

As the PCS1 aggregation pattern at pathogen contact sites was reminiscent of PEN2 localization upon *Bgh* inoculation, we examined the localization of RFP-PCS1 in PEN2-GFP-expressing transgenic plants (Lipka et al., 2005). In pathogen-free plants, RFP-PCS1 showed a strong cytoplasmic fluorescence signal, similar to the GFP-tagged version described above, whereas PEN2-GFP was detectable in the cytoplasm and in the periphery of mobile membrane compartments (Figure S3C-E), recently shown to represent peroxisomes and mitochondria (Fuchs et al., 2016). Pathogen challenge induced colocalization of RFP-PCS1 and PEN2-GFP in aggregate structures with enhanced fluorescence intensity (Figure 2E). RFP-PCS1 did not translocate to all PEN2-GFP-tagged organelles surrounding attempted fungal entry sites, but was confined to a subset with increased PEN2-GFP fluorescence and aggregation (Figure S3E). These organelles were recently identified as a subpopulation of epidermal mitochondria, which become immobile at plant-pathogen contact sites (Fuchs et al., 2016).

To investigate a possible docking role of PEN2 for PCS1, we examined GFP-PCS1 relocalization after inoculation in a *pen2-1* null mutant background (Lipka et al., 2005). GFP-PCS1 is able to aggregate underneath *Bgh* contact sites even in the absence of PEN2 (Figure 2F, S3F), suggesting that mitochondria-associated PEN2 is not necessary for the recruitment of PCS1 to these organelles.

### Phytochelatin synthesis is not required for extracellular defense

When preparing our fusion constructs to investigate the subcellular localization of PCS1 (GFP-PCS1 and PCS1-GFP), we found a striking difference that suggested the two functions of PCS1 in heavy metal tolerance and disease resistance could be uncoupled. Consistent with a previous study (Blum et al., 2010), both GFP fusions were soluble and were found to localize to the cytosol. To verify the functionality of the recombinant proteins, we tested their ability to complement the characteristic penetration resistance and cadmium tolerance defects of the *pen4* mutants. The cadmium hypersensitivity of *pen4* could be complemented by either GFP-PCS1 or PCS1-GFP fusions (Figure 3B), which resulted in both cases in the accumulation of high levels of phytochelatin upon cadmium treatment (Figure 3C), suggesting that both fusions are functional. However, the penetration resistance defect of *pen4* could only be complemented by the GFP-PCS1 fusion (Figure 3A). Similar to GFP-PCS1, RFP-PCS1 was fully functional in both penetration resistance and heavy metal tolerance (Figure S3A-B). Both GFP/RFP-PCS1 and PCS1-GFP relocalize as aggregates at focal sites of attempted fungal penetration (Figure 2A, B), indicating that the lack of complementation of entry resistance in *pen4* by PCS1-GFP is not due to mislocalization of the PCS1-GFP fusion protein. The difference in complementation of the *pen4* disease resistance phenotype by PCS1-GFP versus GFP-PCS1 reveals that the two functions of PCS1 in heavy metal tolerance and penetration resistance can be uncoupled and may therefore reflect an engagement of the protein in two independent pathways. Given that cadmium tolerance has been reported to require phytochelatin synthesis by PCS1 (Grill et al., 1985; Howden et al., 1995; Blum et al., 2007), these results suggest that disease resistance conferred by PCS1 is independent of the ability of the enzyme to synthesize phytochelatin.

**Figure 3:**
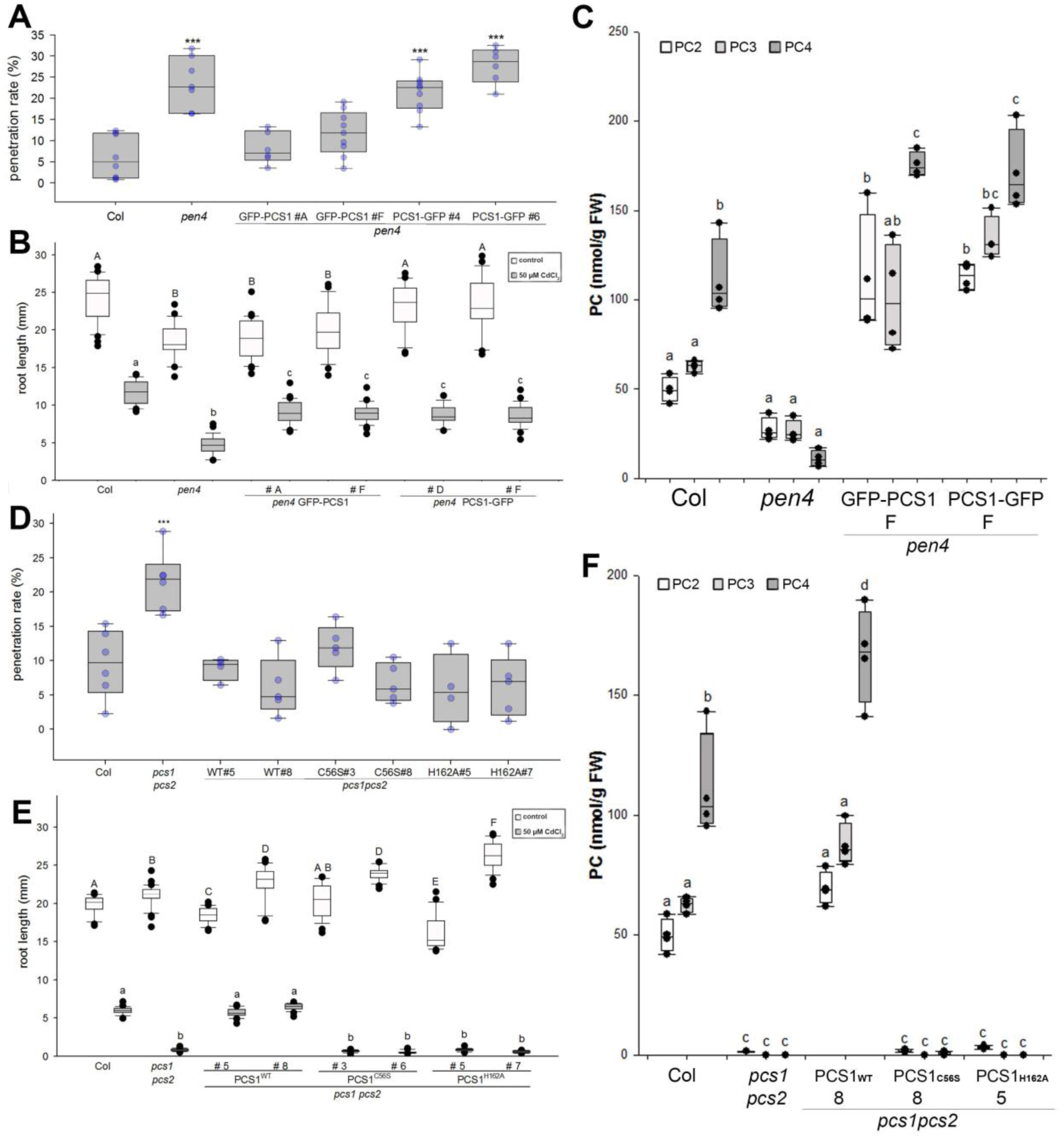
Phytochelatin synthesis is required for heavy metal tolerance but not for non-host resistance. **(A,D)** Box plots showing penetration rate of *Bgh* spores on leaves of 3-week-old Arabidopsis plants. Measurements were done 24 h post-infection on the first true leaves of 5–6 independent plants (600–800 total penetration attempt counted per genotype). *** p<0.001 according to Student’s t-test. **(B, E)** Root length of 7-day-old seedlings grown on with or without 50 µM CdCl_2_. Different letters show statistical differences according to Tukey’s post-hoc test following one-way ANOVA. **(C-F)** Phytochelatin accumulation in 4-day-old seedlings transferred for two days on 50 µM CdCl_2_. Different letters show statistical differences according to Tukey’s post-hoc test following one-way ANOVA.

To test this hypothesis, we generated catalytically inactive PCS1 mutants to investigate the importance of phytochelatin production for fungal resistance. The phytochelatin synthase activity of PCS1 is dependent on a highly conserved catalytic triad located in the N-terminal half of the protein, consisting of Cys56, His162 and Asp180 (Romanyuk et al., 2006). Notably, the cysteine at position 56 has been shown to be involved in the formation of an acyl-enzyme intermediate (Vatamaniuk et al., 2004). Thus, PCS1 proteins carrying C56A, C56S or H162A substitutions lack PC synthesis activity *in vitro* (Vatamaniuk et al., 2004; Romanyuk et al., 2006). To test directly whether the function of PCS1 in fungal resistance is dependent on phytochelatin biosynthesis, we generated transgenic *pcs1pcs2* Arabidopsis plants expressing, under its native promoter, the genomic coding sequence of wild-type PCS1 (PCS1_WT_) or constructs carrying either a C56S (PCS1_C56S_) or H162A (PCS1_H162A_) amino acid substitution in PCS1. As predicted by the inability of the mutated proteins to synthesize phytochelatin (PC) *in vitro* (Vatamaniuk et al., 2004; Romanyuk et al., 2006), transgenic lines expressing PCS1_C56S_ or PCS1_H162A_ were deficient in PC *in vivo* (Figure 3F) and showed the same degree of root growth inhibition as *pcs1pcs2* when grown on 50 µM cadmium chloride (Figure 3E). In contrast, in PCS1_C56S_ or PCS1_H162A_ lines, *Bgh* entry frequencies were similar to those seen in wild-type Col-0 plants or *pcs1pcs2* complemented by PCS1_WT_ (Figure 3D). These results confirm that the function of PCS1 in fungal resistance depends on a PCS1 activity or function that is distinct from phytochelatin synthesis.

### PCS1, but not phytochelatin synthase activity, is involved in indole glucosinolate metabolism to restrict powdery mildew entry in Arabidopsis cells

As suggested by the accumulation of 4-methoxyindol-3-ylmethyl glucosinolate (4MI3G) in *pcs1* plants upon flg22 or Cd^2+^ treatment (Clay et al., 2009; De Benedictis et al., 2018), PCS1, like PEN2, is required for the pathogen-inducible metabolism of IGs. Upon inoculation with a fungal pathogen, *pen2* and *pen4* not only hyperaccumulate 4MI3G (Figure 4B), but additionally are not able to produce indol-3-ylmethylamine (I3A, Figure 4A) and raphanusamic acid (RA, Figure 4D). Levels of IGs and their metabolic products are restored to wild-type levels when the *pen4* or *pcs1pcs2* mutants are complemented by either GFP-PCS1 or the mutated PCS1_C56S_ or PCS1_H162A_, consistent with their wild-type-like disease resistance phenotype (Figure 4B-D). The ABC transporter PEN3/PDR8/ABCG36, although not deficient in I3A or RA (Bednarek et al., 2009), is thought to be involved in this IG metabolic pathway as an exporter of pro-toxins or toxic end products (Stein et al., 2006). This putative role would explain the proper accumulation of I3A/RA and hyper-accumulation of 4-*O*-β-D-glucosyl-indol-3-yl formamide (4OGlcI3F), a product presumably formed upon PEN2-mediated hydrolysis of 4MI3G, in *pen3* leaves inoculated with *Bgh* (Lu et al., 2015). Our analysis of *pcs1* mutants revealed that in addition to I3A and RA, PCS1 is also indispensable for 4OGlcI3F accumulation during the Arabidopsis response to pathogen inoculation (Figure S4A). In addition to 4OGlcI3F hyper-accumulation, some of the *pen3* mutant alleles (e.g. *pen3-1*) exhibit chlorosis after infection with host-adapted mildews (Stein et al., 2006; Lu et al., 2015). This chlorotic phenotype is suppressed in *pen2 pen3-1* (Lu et al., 2015) or *pen3-1 pen4* double mutants (Figure 4E), further supporting the hypothesis that PEN2 and PCS1 act in the same resistance pathway.

**Figure 4.**
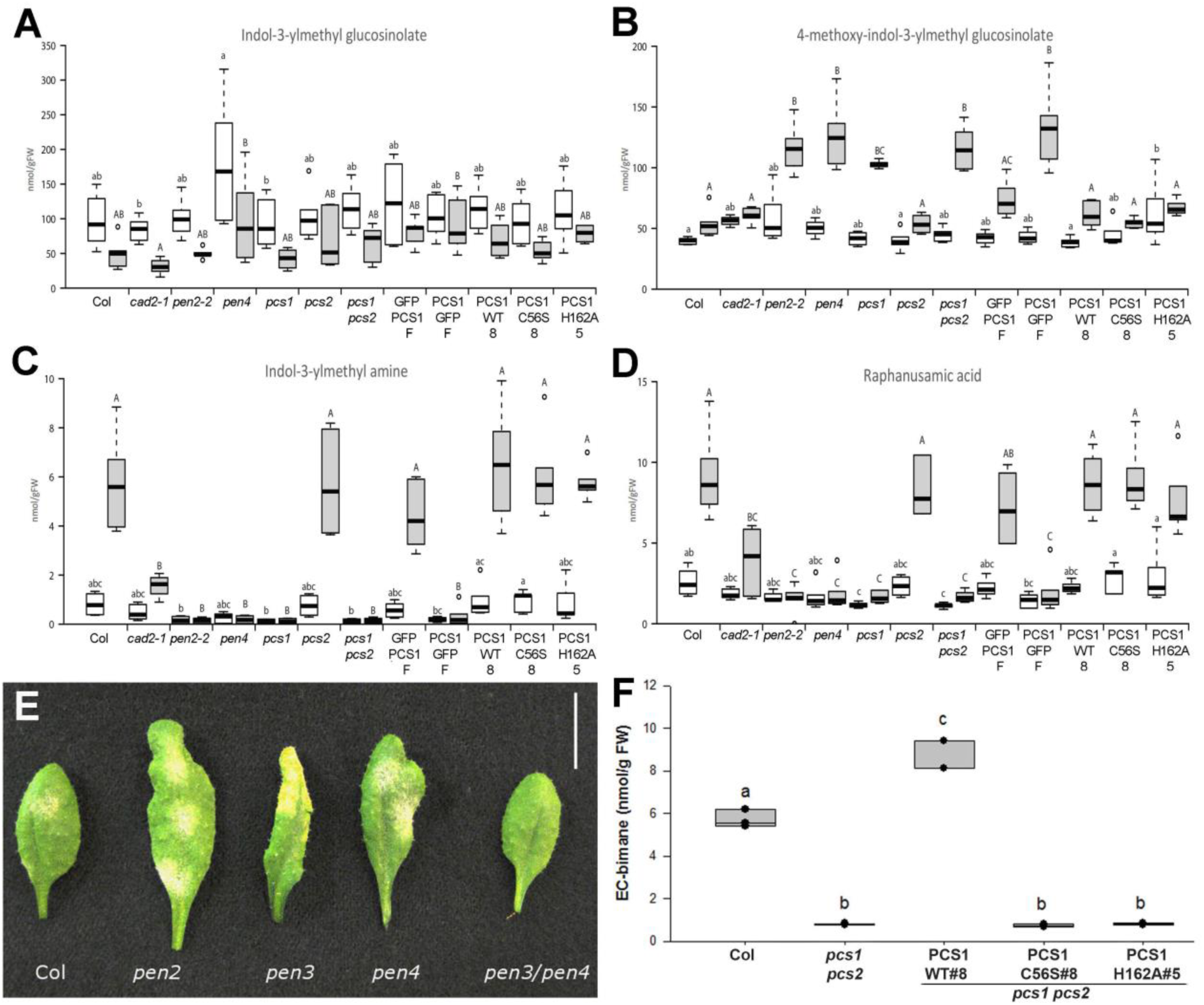
PCS1 is involved in indole glucosinolate metabolism. **(A-D)** Accumulation of selected secondary metabolites, indicated as nmol/g of fresh tissue weight (FW), in Arabidopsis genotypes 16 hours after inoculation with *Bgh* conidiospores. Error bars indicate standard deviations. **(A)** Indol-3-ylmethylglucosinolate (I3G), **(B)** 4-methoxyindol-3-ylmethylglucosinolate (4MI3G), **(C)** Indol-3-ylmethylamine (I3A) **(D)** Raphanusamic acid (RA). Different letters show statistical differences according to Tukey’s post-hoc test following one-way ANOVA. **(E)** Arabidopsis leaves nine days after infection with the adapted mildew *G.cichoracearum* (scale bar is 1 cm). **(F)** EC bimane content in seedling infiltrated with ECG bimane measured by HPLC. Different letters show statistical differences according to Tukey’s post-hoc test following one-way ANOVA.

To directly test whether PEN2 and PEN4 operate in the same pathway, the two mutants were crossed. The *pen2 pen4* double mutant supported a similar increase in entry frequency by *Bgh* as did the *pen2* or *pen4* single mutants (Figure S4). Together with the colocalization with PEN2, these results suggest that PEN2 and PEN4 act in the same pathway conferring extracellular resistance to fungal attack. The correlation between the lack of pathogen-triggered accumulation of I3A and RA and defective entry resistance in *pcs1* lines emphasizes the importance of PCS1 in IG metabolism.

Upon pathogen challenge, the *pen4* mutant hyper-accumulates the physiological substrate of PEN2 myrosinase, 4MI3G, but not its unsubstituted precursor I3G (Figure 4A, B). In contrast, the pathogen-induced accumulation of both IG- and 4MI3G-derived hydrolysis products is essentially abolished (Figure 4C, D). This phenocopies the metabolite profile seen in *pen2* plants (Bednarek et al., 2009; Lu et al., 2015), suggesting that like PEN2, PCS1 may act downstream of 4MI3G biosynthesis to produce I3A, RA and 4OGlcI3F, but upstream of the ABC transporter PEN3/PDR8.

### Is PCS1 peptidase activity required for IG metabolism?

Phytochelatin synthases are known for their γ-glutamylcysteine dipeptidyl transpeptidase activity (EC 2.3.2.15), which catalyzes the transfer of the dipeptide glutamyl-cysteine (EC) from ECG onto another EC unit to generate the polypeptide phytochelatins (EC)_n_-G. In addition, PCS has also been shown to be involved in the turnover of glutathione conjugates of xenobiotics by cleaving the glycine residue of their ECG moiety (Beck et al., 2003; Grzam et al., 2006; Blum et al., 2007).

It has been proposed that the IG-derived ITCs could be glutathionated by GSTU13 and further processed by peptidases including PCS1 (Piślewska-Bednarek et al., 2018). To investigate whether the PC-deficient [*pen4*]-complementing mutants PCS1_C56S_ and PCS1_H162A_ still possessed peptidase activity, we assayed their deglycination activity on the glutathionated bimane (ECG-bimane). None of the PC deficient lines are able to produce EC-bimane from ECG-bimane (Figure 4F). This suggests that, unlike PCS1 metabolism of IG-derived compounds, both PC synthesis and the deglycination of glutathione conjugates depends on the residues C56 and H162 of the catalytic triad.

## DISCUSSION

Phytochelatin synthase is primarily known for its role in tolerance of heavy metals, notably of cadmium (Grill et al., 1989; Clemens et al., 1999). Here, we identified PEN4/*At*PCS1 as a new player in pre-invasive immune responses against fungal pathogens. The second weakly expressed PCS paralogue, *At*PCS2, appears to be dispensable for both heavy metal tolerance and disease resistance. *pcs1* mutants are hypersusceptible to adapted and non-adapted fungi, unveiling the importance of PCS1 for defense against fungal invasion. This impaired pre-invasive immunity might also explain the previously reported strong cell death phenotype in *A. thaliana pcs1* plants in response to inoculation with the non-adapted oomycete pathogen *Phytophthora infestans*, because invasive growth of this oomycete in *pen2* plants is also linked to a localized plant cell death response (Lipka et al., 2005; Kühnlenz et al., 2015). Like the other PEN proteins, PCS1 concentrates underneath the penetration site of fungal appressoria. Interestingly, in uninfected cells PCS1 is cytosolic, and in response to infection it relocalizes to an endomembrane compartment(s) at incipient entry sites of attack. We did not observe this aggregation phenomenon after cadmium treatment, further underlining the dual functions of PCS1. Colocalization of RFP-PCS1 and PEN2-GFP in plant cells under attack suggests that these compartments are mitochondria as previously reported (Fuchs et al., 2016). Recent proteome studies localized PCS1 to the cytosol (Ito et al., 2011) and to the *A. thaliana* mitochondrial complexome (Senkler et al., 2017), supporting its dual localization capacity.

Unlike the *pen1 pen2* double mutant (Lipka et al., 2005), the susceptibility of the *pen2 pen4* mutant to fungal infection is not more pronounced than either of the single mutants (Figure S4). This suggests that PEN2 and PEN4/PCS1 act in the same pathway. PEN2 has been shown to be an essential component of an IG metabolism pathway involved in broad-spectrum resistance against filamentous eukaryotic pathogens (Bednarek et al., 2009; Pastorczyk and Bednarek, 2016). PCS1 appears to be equally involved in this pathway as the *pcs1* mutants exhibit exactly the same I3A, RA and 4OGlcI3F deficiency as *pen2* plants (Figure 4A,B and S4A). The colocalization of both enzymes in response to pathogen attack could facilitate stimulus-induced metabolite channelling of these metabolites, which in turn could account for the lack of detection of the intermediates between I3G and I3A/RA or 4MI3G and 4OGlcI3F. However, this does not allow us to unambiguously determine whether PEN4 is upstream or downstream of PEN2.

The shared phenotype of the *pen2* and *pen4* mutants and the colocalization of PEN2 and PEN4 in the same compartment at the site of fungal ingress suggested a possible protein-protein interaction. However, we did not obtain any positive interactions between PEN2 and PEN4 in split-ubiquitin yeast two-hybrid experiments (Figure S5A), nor could we co-immunoprecipitate these two proteins from infected leaves (Figure S5B). The lack of physical interaction is confirmed by the fact that PCS1 is still able to relocalize in a null *pen2* mutant background (Figure 4F and S3F). Focal accumulation of the PEN3 ABC transporter as disks in the plasma membrane and colocalization of PEN2 and PEN4 aggregates on mitochondria underneath pathogen contact sites rather points to pathogen-inducible macromolecular crowding as a potential alternative mechanism. Such an arrangement of PEN2 and PEN4 would facilitate metabolite channelling for targeted release of antimicrobials (Ellis, 2001; Stein et al., 2006; Fuchs et al., 2016; Guigas and Weiss, 2016). However, it remains to be tested whether enforced PEN4 mislocalization impairs its activity in extracellular defense.

Using mutated versions of PCS1, we showed that the function of PCS1 in IG metabolism does not require phytochelatin synthase activity *sensu stricto* as we could generate PC-deficient mutants functional for IG metabolism. Indeed the ability to metabolize IGs is retained in mutant versions of PCS1 (Figure 4A-D) that cannot produce any phytochelatins (Figure 3F) nor cleave the ECG bimane conjugate (Figure 4F). Our findings confirm and extend a previous study in which heterologous expression of *Caenorhabditis elegans PCS* in the Arabidopsis *pcs1* background complemented cadmium hypersensitivity, but not a leaf cell death phenotype in response to *P. infestans* inoculation (Kühnlenz et al., 2015). Residue Cys56 of the catalytic triad (Cys56, H162, Asp180) has been implicated in the formation of an acyl-enzyme intermediate with γ-EC. There appears to be an as yet unidentified second acylation site in the C-terminal part of PCS1, which shows a low level of sequence conservation across plant species (Vatamaniuk et al., 2004; Rea, 2012). This second site, which is absent in prokaryotic PCS, could potentially participate in IG metabolism. This hypothesis is supported by the fact that the PCS from *C. elegans* that was not able to complement the cell death phenotype in *pcs1* plants has a significantly shorter C-terminal part compared to *At*PCS1 (Rea et al., 2004; Kühnlenz et al., 2015). This assumed significance of the PCS1 C-terminus could also explain why GFP-PCS1, but not the PCS1-GFP fusion, is non-functional in IG metabolism and extracellular defense despite its proper subcellular localization. Alternatively, PCS1 may not act catalytically but rather it may stabilize a chemically labile intermediate in the IG metabolism pathway or stabilize a protein complex required for IG catabolism (Figure S6).

The chemical structures of RA and I3A, combined with the fact that their formation is dependent on ECG supply via γ-glutamylcysteine synthetase, suggested indol-3-ylmethyl-ITC-ECG conjugates as intermediates in pathogen-triggered IG metabolism in Brassicaceae (Bednarek et al., 2009). The existence of such conjugates *in planta* is supported by the identification of indolyl-dithiocarbamate glucoside during engineering of brassinin (an IG-derived phytoalexin from *Brassica* spp.) biosynthesis in *Nicotiana benthamiana* (Klein and Sattely, 2017). GSTU13 was recently found to be essential for pathogen-inducible formation of RA, I3A, and 4OGlcI3F, suggesting that it catalyzes *in planta* the conjugation of unstable glucosinolate hydrolysis products, the indol-3-ylmethyl-ITCs, with ECG to form a dithiocarbamate adduct (Piślewska-Bednarek et al., 2018). Although the exact mechanism of PCS1 in this pathway remains to be defined, our combined genetic and metabolite analysis identifies PCS1 as plausible candidate for further processing of this adduct to I3A and RA, and the corresponding 4-methoxy adduct to 4OGlcI3F. In conclusion, AtPCS1 is a multi-functional protein: it detoxifies xenobiotics via chelation of heavy metals by phytochelatin or via degradation of glutathione-conjugates, and it participates in the production of antimicrobials through the processing of IG-derived compounds.

## MATERIAL AND METHODS

### Plant and fungal lines and growth conditions

*Arabidopsis thaliana* plants were grown 15 to 20 days in growth chambers at 22°C with a 12-h photoperiod. All Arabidopsis mutants and transgenic lines were in the Columbia background. Host powdery mildew (*Golovinomyces cichoracearum* UCSC1) was cultured on squash for 10 to 12 days, and then applied to Arabidopsis using settling towers. Non-host barley powdery mildew (*Blumeria graminis f. sp hordei* CR3) was grown on barley (*Hordeum vulgare*) line Algerian-S (CI-16138) and inoculated onto Arabidopsis using the methods described by Zimmerli et al. (2004).

Transgenic Arabidopsis lines were generated according to the floral dip method (Clough and Bent, 1998), T1 lines carrying single T-DNA insertion events were identified by screening their progeny for segregation of antibiotic resistance in the appropriate ratio. Resistant T2 seedlings were transferred to soil and used in pathogen assays, in which several representative independent lines were selected. T2 individuals homozygous for T-DNA insertions (based on 100% antibiotic resistance or 100% GFP fluorescence among their T3 progeny) were further used for pathogen resistance or heavy metal tolerance assays and metabolic analysis.

For assaying tolerance to cadmium, Arabidopsis seeds were surface sterilized and sown onto agar plates containing half-strength Murashige and Skoog medium and 50 µM cadmium chloride (CdCl_2_). Plates were grown under continuous light at 22 °C. After 7 days, plates were scanned and root lengths measured from scanned images using the software ImageJ. All experiments were repeated at least three times with similar results.

### Map-based cloning of PEN4

The *pen4* mutation was mapped to the bottom arm of chromosome 5 using an F2 mapping population generated from a cross between *pen4* and Landsberg-*erecta* (Ler), according to Lukowitz et al. (2000). We narrowed down the region containing *PEN4* to a genomic interval of approximately 20 kbp at the end of BAC clone MRH10. Analysis of lists of genes that co-express with either *PEN1*, *PEN2* or *PEN3* across publicly available microarray datasets identified a gene highly correlated in expression with all three PEN genes (Fig.S2) and that also resided in the 20 kbp mapping interval. Sequencing of this gene, At5g44070, from the *pen4* mutant showed that it harbored a G to A transition at nucleotide 1713 from the A of the translational start site of the genomic sequence. This mutation results in a predicted truncation of the AtPCS1 protein at residue 236.

### Plasmids and DNA constructs

PCS1 fusions to GFP or RFP were constructed using Gateway technology (Invitrogen). The Arabidopsis PCS1 (AtPCS1) genomic coding sequence was amplified by PCR using BAC clone MRH10 as template DNA and recombined into donor vector pDONR/Zeo (Invitrogen). The resulting entry vector was then recombined by gateway LR reaction with the destination vectors pMDC43 and pMDC83 (Curtis and Grossniklaus, 2003) to produce expression clones GFP-PCS1 and PCS1-GFP, respectively. pDONR/Zeo AtPCS1 was recombined with a modified pEG106 (Gutierrez et al. 2009) containing an HA-tagged mCherry to generate RFP-PCS1. GFP fusions were transformed into a *pen4* mutant background (and selected on hygromycine 50 mg/L) and RFP-PCS1 into a *pcs1pcs2* double mutant (Blum et al. 2007, selected on soil after BASTA spray). For complementation analyses, the entire PCS1 genomic coding sequence in addition to 3 kbp of promoter sequence upstream of the translational start site and 1 kbp downstream of the translational stop codon was amplified by PCR using BAC clone MRH10 as template DNA and recombined into donor vector pDONR/Zeo.. The Quickchange mutagenesis kit (Stratagene) was used to introduce mutations in pDONR/Zeo PCS1_WT_, resulting in a change of cysteine 56 to serine (PCS1_C56S_) or of histidine 162 to alanine (PCS1_H162A_). The resulting entry vectors were then recombined with destination vector pGWB1 (conferring resistance to hygromycin) (Nakagawa et al. 2007) to generate expression clone PCS1-WT These mutated PCS1 genomic clones were further recombined in the binary vector pGWB1. All expression clones were introduced into *Agrobacterium tumefaciens* (strain GV3101) and the resulting strains were used for stable transformation of Arabidopsis according to the floral dip method.

### Generation of double mutants

Double mutants were generated by crossing the *pen4* mutant to mutants *pen2-1*, *pen2-3* or *pen3-1*. F2 individuals homozygous at the mutated loci were identified using cleaved amplified polymorphic sequence (CAPS) markers described. For the CAPS marker to identify the *pen4* mutation, the *pen4* sequence was PCR amplified with forward primer 5’taagacaaaacttgtggattgg3’ and reverse primer 5’cacccatctgatgaattgg3’ and the resulting DNA product cleaved using the *Mbo*I restriction enzyme.

### Microscopic observations

*Bgh* penetration assays were carried out as described by Zimmerli et al. 2004), and stained with aniline blue according to Vogel et al. 2000. Successful penetrations (forming haustoria with large callose encasement) and failed penetrations (forming callose papillae) were counted to determined the penetration rate (succesfull penetration / number of attempts). For imaging of fluorescent proteins in Arabidopsis epidermal cells, leaves from 2 to 3-week-old plants were mounted in a 0.01 mg/mL propidium iodide solution in water (to stain fungal structures). Leaves were examined between 12 and 18 hours following fungal inoculation on spinning disc confocal microscope consisting of a Leica DMI 6000 B inverted microscope (Leica Microsystems, Wetzlar, Germany) fitted with a Yokogawa CSU-10 spinning disc confocal attachment (Yokogawa Electric Corporation, Tokyo, Japan) and a Photometrics QuantEM 512SC EM-CCD camera (Photometrics, Tucson, AR). Samples were mounted in water and observed with a 63x water immersion objective. EGFP was excited at 488 nm, and fluorescence was collected through a 525/50 nm band-pass filter and fungal structures stained with propidium idodide at 620/60 band-pass filter (Chroma Technologies, Brattleboro, VT). RFP (mCherry) was excited at 561 nm, and fluorescence was collected through a 620/60 nm band-pass filter (Chroma Technologies, Brattleboro, VT). Microscope control and acquisition of images and z-series were accomplished using Metamorph software (Molecular Devices, Sunnyvale, CA) and image processing was performed by using ImageJ (W. Rasband, National Institutes of Health, Bethesda, MD) software.

### Metabolic profiling

Analysis for phytochelatin biosynthesis were performed according to Blum et al. (2007). Extraction and analysis of indole glucosinolates and their derivatives were performed according to Bednarek et al. (2009) and Lu et al. (2015).

## AUTHOR CONTRIBUTIONS

KH, ML, VL, AM, EG, PSL and SS designed the research, KH, ML, CC, PB, MPB, CSR, MS, RF, CK performed research and analyzed data. KH and SS wrote the manuscript with contributions from PB, VL and PSL

## ACKNOWLEDGEMENTS

We acknowledge the work of the late Ralph Blum for metabolite analysis and data processing shown in Figure 3C, F and Figure 4F.

We are grateful to Chris Cobbett (University of Melbourne, Australia) for providing the *cad1* mutants and to Phil Rea (University of Philadelphia, USA) for AtPCS1 cDNA; Ryan Guttierez (Carnegie Institution of Science, Stanford, USA) for the pEG106 mCherry vector; Chris Cobbett, Phil Rea and Stephan Clemens (IGB, Jena, Germany) for helpful discussions; Bill Underwood (Energy Biosciences Intitute, Berkeley, USA) for his advice and help in microscopy and the Somerville lab for stimulating discussion.

Financial support was provided by the National Science Foundation and the Carnegie Institution of Science (to SS), a Stanford Graduate Fellowship (to MS), the Max Planck Society (to PSL), by Deutsche Deutsche Forschungsgemeinschaft (DFG) Research Grant SPP1212 (to PSL), DFG grant GR 938 (to EG) and DFG grant LI 1317/2-1 (to VL), by Spanish Ministry of Economy and Competitiveness (MINECO) grants BIO2015-64077-R and BIO2012-32910 (to AM) and the National Science Centre grant 2012/07/E/NZ2/04098 (to PB).

## SUPPORTING INFORMATION

Table S1: List of primers used in this study

**Fig. S1. *pen4* is allelic to *cad1*.**

Box plots showing penetration rate of *Bgh* spores on leaves of 3-week-old Arabidopsis plants. Measurements were done 24 h post-infection on the first true leaves of 5–6 independent plants (600–800 total penetration attempt counted per genotype). *** p<0.001; * p<0.05 according to Student’s t-test.

**Fig. S2. *pen4* is more susceptible to various fungal pathogens**

Quantification of infection by *Plectosphaerella cucumerina* (A) or *Botrytis cinerea* (B, 7dpi) and the host biotrophic mildew *Golovinomyces cichoracearum* (C, 7dpi). (**A**) Percentage of decayed plants of the indicated genotypes at different days after inoculation (dpi) with at the rate of 4 × 10^6^ *P. cucumerina* spores/ml. Data represent the average (±SE) from one of three independent experiments that gave similar results. (B) Percentage of plant fresh weight (FW) reduction of the indicated genotypes at 7 days after inoculation with at the rate of 5 × 10^4^ *B. cinerea* spores/ml. Data represent the average (±SE) of three independent experiments. (**C**)

**Fig. S3: Upon pathogen attack, the functional fusion protein RFP-PCS1 translocates from the cytosol to the surface of immobilized mitochondria tagged withPEN2-GFP.**

(A-B) RFP-PCS1 fusion complements *pcs1pcs2* mutant phenotypes. (A): penetration rate of *Blumeria graminis* on *Arabidopsis thaliana* leaf epidermal cells. (B) root growth inhibition of 7-day-old seedling on 50 µM CdCl_2_.

(C-E) Confocal images show unchallenged (C) and challenged (D, E) leaf epidermal cells of stable double transgenic plants transformed with P_*PEN2*_::PEN2-GFP and P_*35S*_::RFP-PCS1 constructs. In unchallenged epidermal cells (C), RFP-PCS1 shows a cytoplasmic localization pattern, whereas PEN2-GFP can be detected in the cytoplasm and the periphery of mobile membrane compartments (recently demonstrated to represent mitochondria and peroxisomes). Pathogen challenge triggers colocalization patterns in aggregate structures with enhanced fluorescence intensity (D,E). Notably, in contrast to PEN2-GFP, RFP-PCS1 cannot be detected in the periphery of organelles (E, white arrows). 20 hpi with *Bgh*.

(F) Transgenic *pen2-1* knockout lines expressing RFP-PCS1 show pathogen-induced aggregate formation in absence of PEN2. 19 hpi with *Bgh.* White dotted line indicates position of the fungal appressorium. ap, appressorial germ tube. Bars = 10 µm.

**Fig. S4. PEN2 and PCS1 act in the same biochemical and immune pathways**

A) Accumulation of 4-*O*-β-D-glucosyl-indol-3-yl formamide, indicated as nmol/g of fresh tissue weight (FW), in different Arabidopsis genotypes 16 hours after inoculation with *Bgh* conidiospores. (B) Penetration rate of *pen2 pen4* double mutant by *Bgh*. Different letters show statistical differences according to Tukey’s post-hoc test following one-way ANOVA.

**Fig. S5. PCS1-PEN2 physical interaction assays.**

(A) Split Ubiquitin Y2H assays suggest no direct interaction of PEN2 and PCS1. Only yeast cells containing the positive control KAT1 grow on selective medium (-Ade, -His).

(B) Co-immunoprecipitation experiments suggest that PEN2 and PCS1 do not interact with each other. 3-week-old plants expressing PEN2-GFP and 3xHA-RFP-PCS1 were infected (Inf) or not (UI). Total leaf protein extracts were subjected to a PEN2-immunoprecipitation (anti-GFP agarose beads) and purified fractions (B: bound, UB: unbound) were tested by western blotting against PEN2 (anti-GFP) and PCS1 (anti-HA).

**Fig. S6. Function of PCS1 in pathogen-triggered indole glucosinolate metabolism.**

As indicated by a deficiency in indole glucosinolate hydrolysis products and hyper-accumulation of 4-methoxy-indol-3-ylmethyl glucosinolate in *pcs1* mutant plants, PCS1 acts downstream of intact glucosinolates. PCS1 might contribute to the processing of the isothiocyanate-glutathione adduct or modulates the activity of other enzymes of the pathway.

---

**Table 1.**
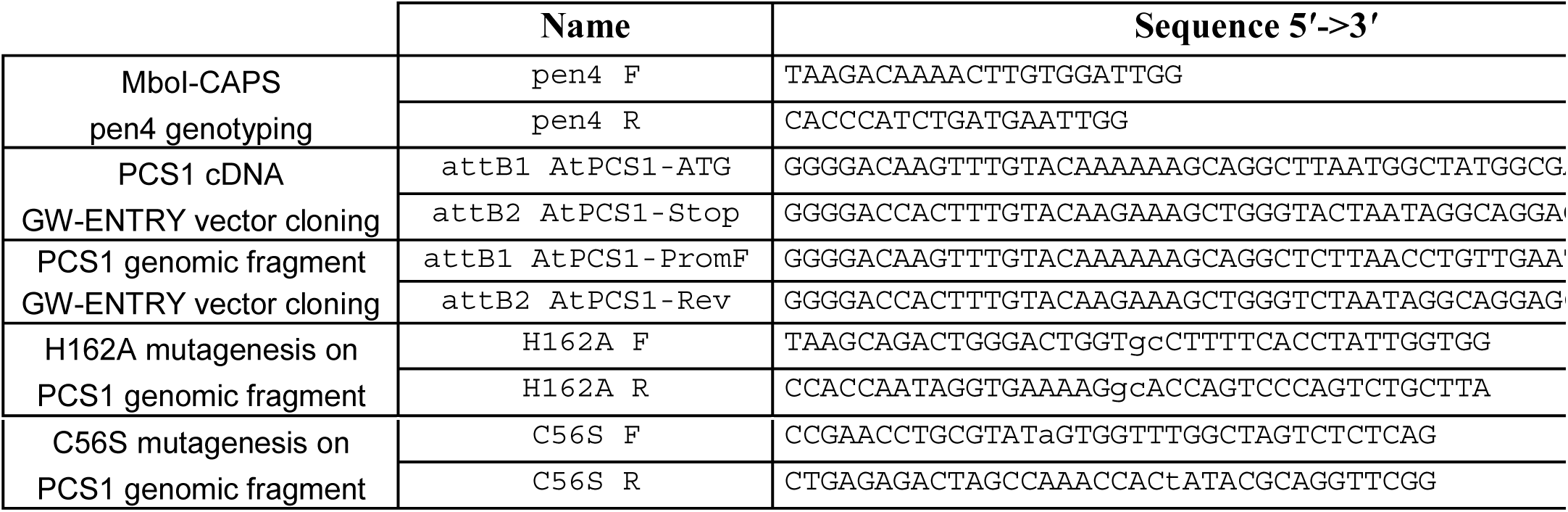
Primers used in this study

