## Supplemental Figures for "Arabidopsis phytochelatin synthase 1, but not phytochelatin synthesis, functions in extracellular defense against multiple fungal pathogens"

**FIGURE S1**

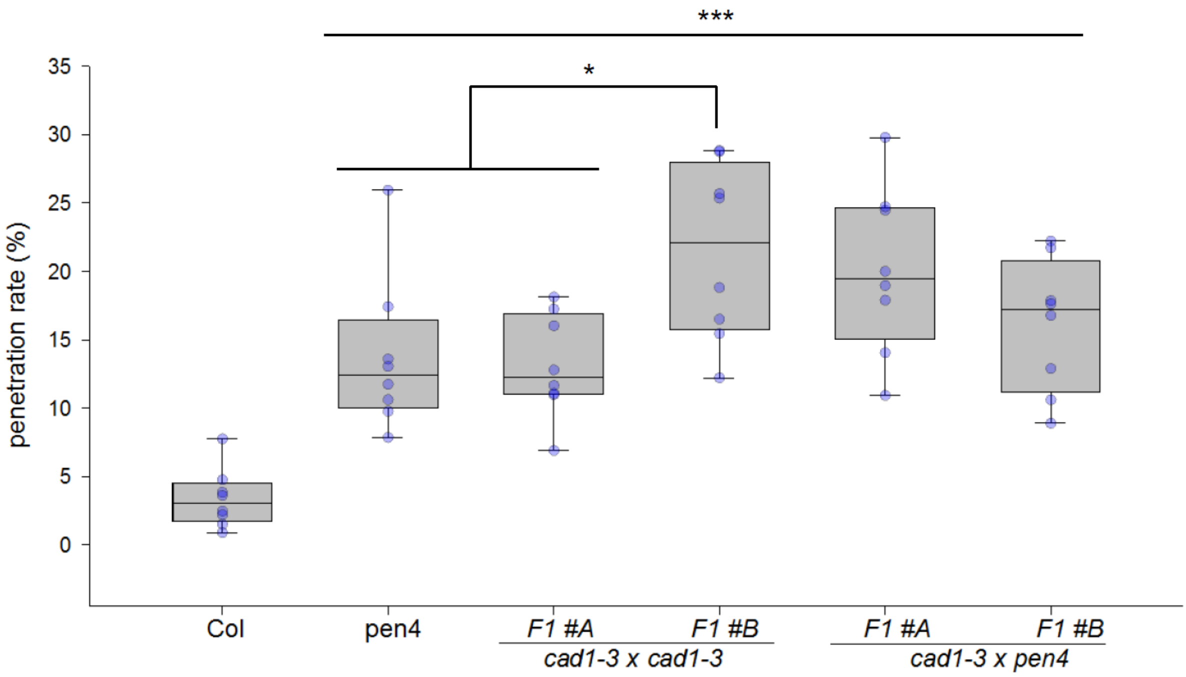


**Fig. S1. *pen4* is allelic to *cad1/pcs1*.**

Box plots showing penetration rate of *Bgh* spores on leaves of 3-week-old Arabidopsis plants. Measurements were done 24 h post-infection on the first true leaves of 5–6 independent plants (600–800 total penetration attempt counted per genotype). *** p<0.001; * p<0.05 according to Student's t-test.

**FIGURE S2**

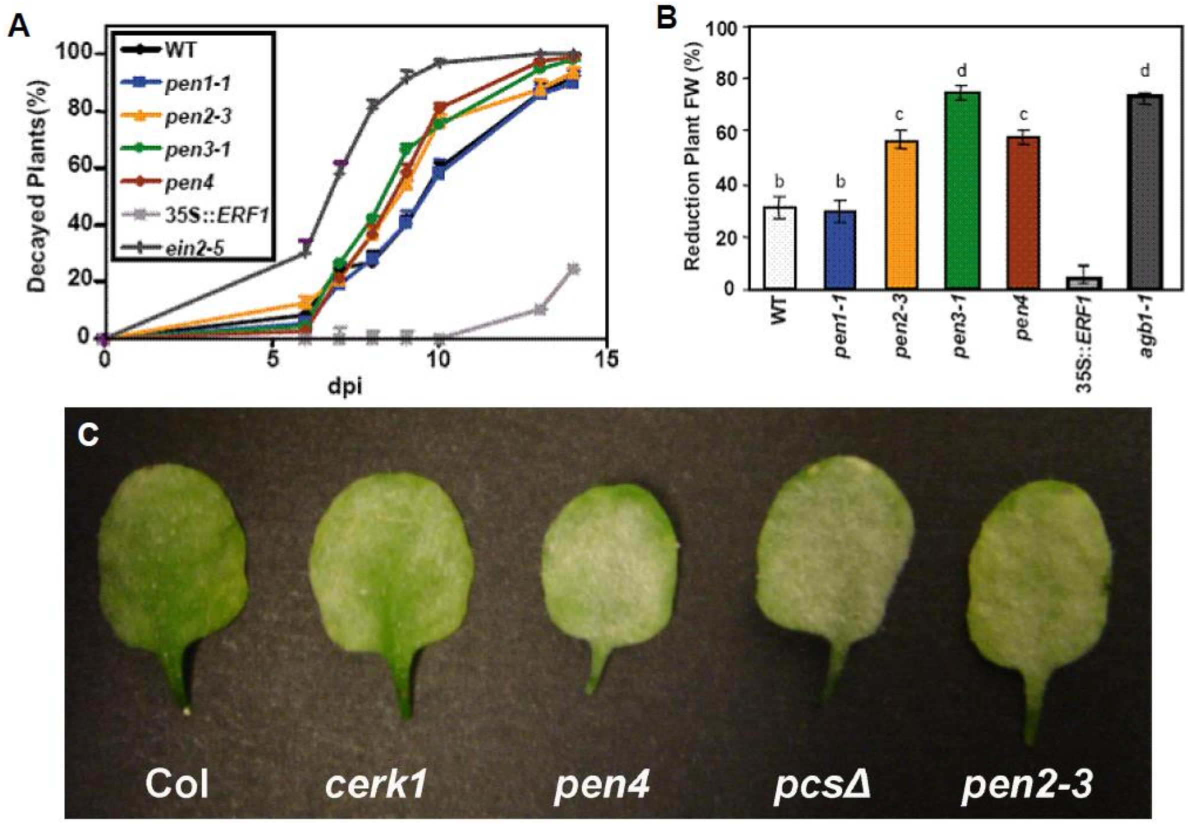


**Fig. S2.** ***pen4* is more susceptible to various fungal pathogens**

Quantification of infection by *Plectosphaerella cucumerina* (A) or *Botrytis cinerea* (B, 7dpi) and the host biotrophic mildew *Golovinomyces cichoracearum* (C, 7dpi). (**A**) Percentage of decayed plants of the indicated genotypes at different days after inoculation (dpi) with at the rate of $4 \times 10^6$ *P. cucumerina* spores/ml. Data represent the average (±SE) from one of three independent experiments that gave similar results. (B) Percentage of plant fresh weight (FW) reduction of the indicated genotypes at 7 days after inoculation with at the rate of $5 \times 10^4$ *B. cinerea* spores/ml. Data represent the average (±SE) of three independent experiments.

### FIGURES S3

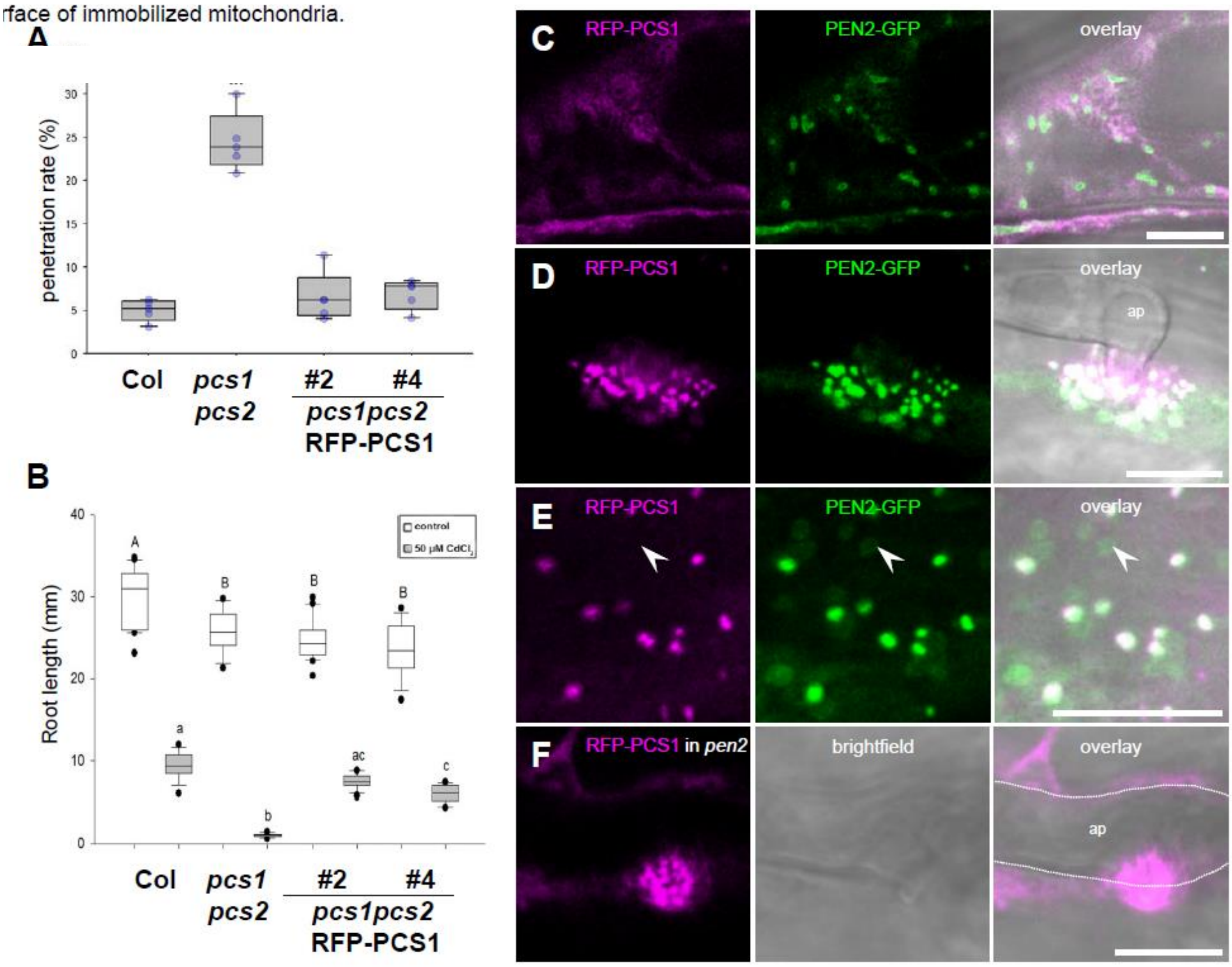


**Fig. S3: Upon pathogen attack, the functional fusion protein RFP-PCS1 translocates from the cytosol to the surface of immobilized mitochondria tagged withPEN2-GFP.**

(A-B) RFP-PCS1 fusion complements *pcs1pcs2* mutant phenotypes. **(**A): penetration rate of *Blumeria graminis* on *Arabidopsis thaliana* leaf epidermal cells. (B) root growth inhibition of 7-day-old seedling on 50 µM $CdCl_2$. (C-E) Confocal images show unchallenged (C) and challenged (D, E) leaf epidermal cells of stable double transgenic plants transformed with $P_{PEN2}$::PEN2-GFP and $P_{35S}$::RFP-PCS1 constructs. In unchallenged epidermal cells (C), RFP-PCS1 shows a cytoplasmic localization pattern, whereas PEN2-GFP can be detected in the cytoplasm and the periphery of mobile membrane compartments (recently demonstrated to represent mitochondria and peroxisomes). Pathogen challenge triggers co-localization patterns in aggregate structures with enhanced fluorescence intensity (D,E). Notably, in contrast to PEN2-GFP, RFP-PCS1 cannot be detected in the periphery of organelles (E, white arrows). 20 hpi with *Bgh*. (F) Transgenic *pen2-1* knockout lines expressing RFP-PCS1 show pathogen-induced aggregate formation in absence of PEN2. 19 hpi with *Bgh.* White dotted line indicates position of the fungal appressorium. ap, appressorial germ tube. Bars = 10 µm.

**FIGURE S4**

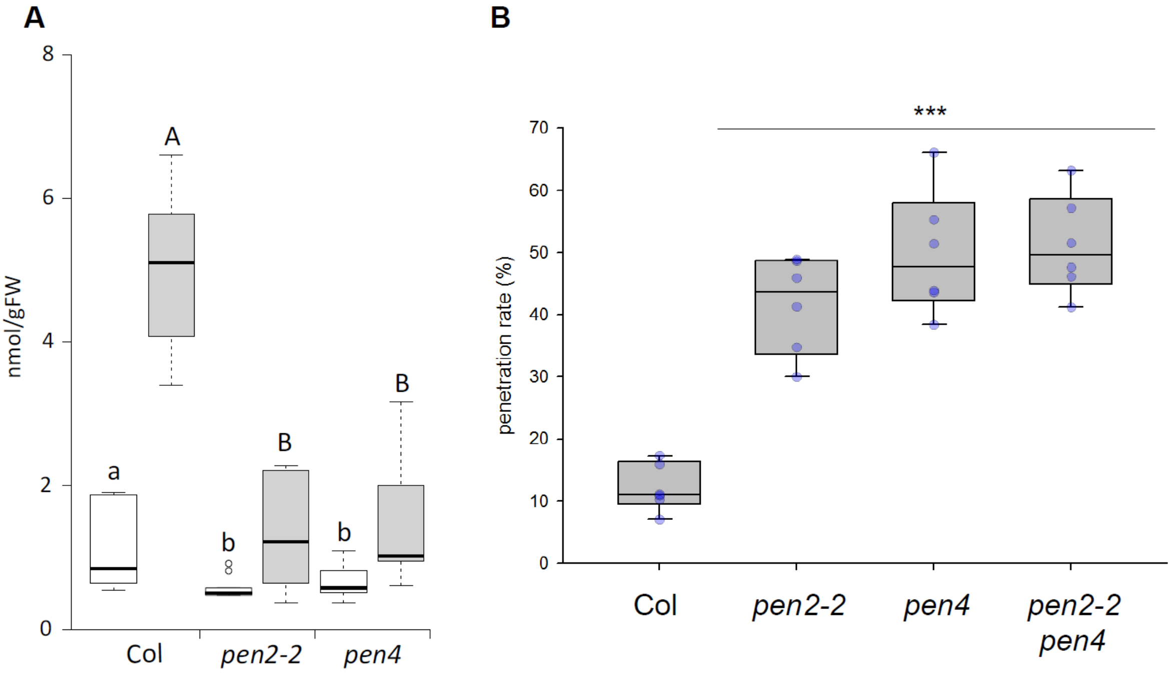


**Fig. S4. PEN2 and PCS1 act in the same biochemical and immune pathways**

(A) Accumulation of 4-*O*-β-D-glucosyl-indol-3-yl formamide, indicated as nmol/g of fresh tissue weight (FW), in different Arabidopsis genotypes 16 hours after inoculation with *Bgh* conidiospores. (B) Penetration rate of *pen2 pen4* double mutant by *Bgh*. Different letters show statistical differences according to Tukey's post-hoc test following one-way ANOVA.

**FIGURE S5**

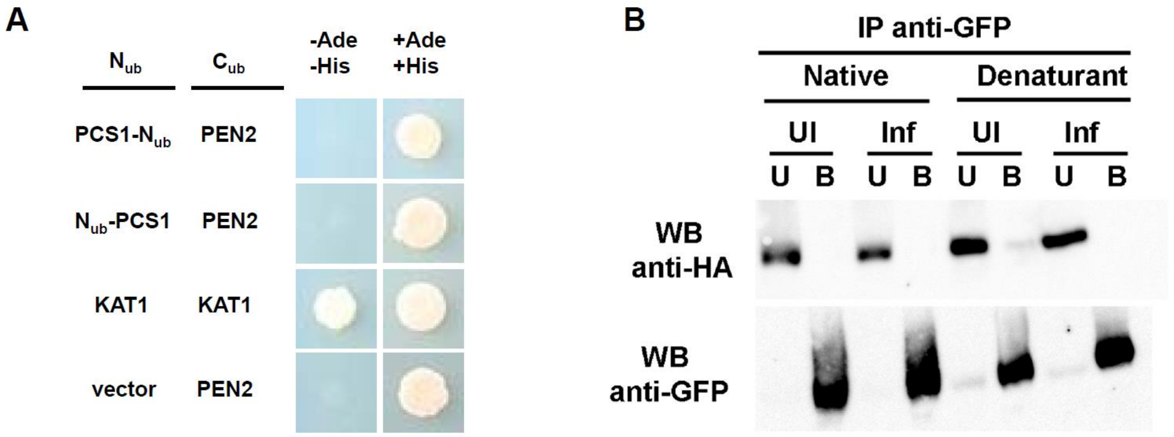


**Fig. S5. PCS1-PEN2 physical interaction assays.**

(A) Split Ubiquitin Y2H assays suggest no direct interaction of PEN2 and PCS1. Only yeast cells containing the positive control KAT1 grow on selective medium (-Ade, -His).

(B) Co-immunoprecipitation experiments suggest that PEN2 and PCS1 do not interact with each other. 3-week-old plants expressing PEN2-GFP and 3xHA-RFP-PCS1 were infected (Inf) or not (UI). Total leaf protein extracts were subjected to a PEN2-immunoprecipitation (anti-GFP agarose beads) and purified fractions (B: bound, UB: unbound) were tested by western blotting against PEN2 (anti-GFP) and PCS1 (anti-HA).

**FIGURE S6**

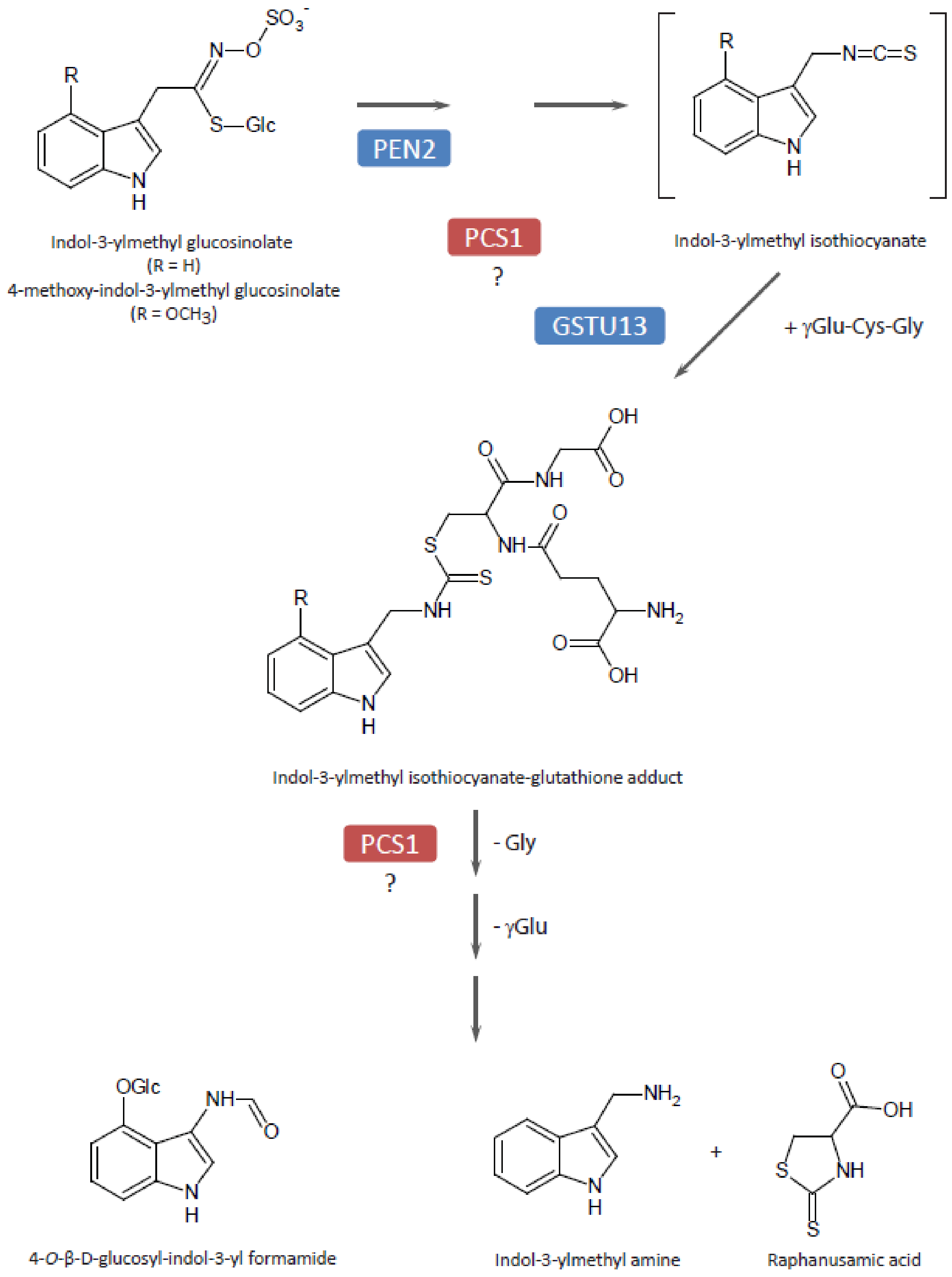


**Fig. S6. Function of PCS1 in pathogen-triggered indole glucosinolate metabolism.**

As indicated by a deficiency in indole glucosinolate hydrolysis products and hyper-accumulation of 4-methoxy-indol-3-ylmethyl glucosinolate in *pcs1* mutant plants, PCS1 acts downstream of intact glucosinolates. PCS1 might contribute to the processing of the isothiocyanate-glutathione adduct or modulates the activity of other enzymes of the pathway.
